# Design of a Streptolysin O Epitope-Centric Nanoparticle Vaccine Against *Streptococcus pyogenes*

**DOI:** 10.1101/2025.05.25.655993

**Authors:** Di Tang, Yashuan Chao, Elisabeth Hjortswang, Joel Ströbaek, Lucas Hultgren, Tirthankar Mohanty, Christofer Karlsson, Simon Ekström, Lotta Happonen, Oonagh Shannon, Lars Malmström, Johan Malmström

## Abstract

*Streptococcus pyogenes* (Group A *Streptococcus*, GAS) is a significant human pathogen for which no licensed vaccine is currently available. Here, we report a *de novo* designed epitope-centric protein-based nanoparticle vaccine against GAS. By integrating structural mass spectrometry techniques and deep learning approaches, we re-engineered a protective epitope (D3m) present in domain 3 of streptolysin O, a prominent pore-forming toxin produced by GAS. D3m was displayed on the surface of a self-assembling icosahedral nanoparticle (D3m-NP) to enhance epitope presentation and immunogenicity. Mice immunised with D3m-NP mounted haemolysis-neutralising titres and displayed a more uniform, epitope-centric antibody response than those receiving the community-standard detoxified full-length streptolysin O. Our findings highlight a promising strategy for GAS vaccine development by combining multimodal protein mass spectrometry, protein design and a versatile protein-based nanoparticle vaccine platform.

## Introduction

Group A *Streptococcus* (GAS), or *Streptococcus pyogenes*, is a strict human pathogen linked to a diverse spectrum of diseases, making it a persistent and significant threat to global public health ^1^. Clinical manifestations range from mild illnesses such as pharyngitis to severe invasive conditions including necrotising fasciitis and sepsis ^2^. Furthermore, post-infectious sequelae such as rhematic heart disease contribute to long-term morbidity and impose a substantial burden on healthcare systems ^1^. GAS produces a variety of surface-associated and secreted proteins that exert multifunctional effects to modulate and evade the host immune system. Several of these proteins are essential for bacterial pathogenesis, however, the complexity also poses a significant challenge for the development of an effective vaccine ^3^. Despite considerable efforts over the past decades, no safe and effective vaccine has been approved for GAS although several important streptococcal virulence factors, *e.g.*, M-protein, C5a peptidase and streptolysin O, have been shown to induce protective immunity in animal models ^4^.

Streptolysin O (SLO) is a 60 kDa pore-forming exotoxin secreted by GAS. The mature form of SLO begins with a 69-residue intrinsically disordered region at the N-terminus, followed by three non-contiguous domains (domain 1-3), and ends with a C-terminal membrane-binding domain 4 ^5^. SLO primarily targets cholesterol-rich cellular membranes, where it oligomerises to form a prepore complex. The oligomerisation triggers a drastic conformational change initiated by domain 3 that enables membrane penetration and pore formation, resulting in cytolysis and cell death ^6,7^. In addition to the cytolytic activity, SLO contributes to GAS pathogenesis through multiple pathomechanisms such as translocating co-toxin NAD+ glycohydrolase into host cells ^8^, enhancing streptococcal superantigen activity ^9^, modulating neutrophil responses ^10,11^, and impairing both phagocytic clearance ^12,13^ and intracellular lysosomal killing of GAS ^14^. SLO is a highly conserved cytolysin present in nearly all GAS strains and is a key target in several multicomponent subunit vaccines currently in pre-clinical development ^15,16^. Both recombinant protein and lipid-based mRNA forms of SLO immunogens have been tested, but to date there is no licensed vaccine ^17,18^. Current formulations of SLO-based protein subunit vaccines typically introduce point mutations in domain 4 to generate a full-length detoxified construct, which prevents SLO binding to cellular membranes ^19^. However, the introduction of point mutations may disrupt important native epitopes that can compromise the protective efficacy of the elicited antibody responses *in vivo*. A broader limitation of current full-length protein subunit approaches is the immunodominance hierarchy, which drives the immune responses toward immunodominant but potentially non-functional or weakly protective epitopes ^20^. Furthermore, combining several different antigens into multicomponent protein subunit vaccines puts high demands on finding the optimal conditions to maintain the solubility and to ensure the desirable solution-state conformation of all included antigen components. These challenges underscore the importance of exploring new vaccine platforms to generate stable immunogens that can present several protective epitopes in their correct three-dimensional conformation, which is critical to ensure the development of stable and effective vaccines ^21^.

Recently, computationally designed hyper-stable protein-based nanoparticles have emerged as a promising approach to precisely modulate selection, valency, presentation, and spacing of target antigens ^22–27^. These protein nanoparticles assemble from two components to form multi-subunit symmetric protein nanostructures with molecular weight of up to 2.8 megadalton and diameters of 24-40 nm ^28^. The ability to precisely engineer antigen or epitope structures on designed nanoparticles offers a powerful platform for multivalent presentation *in vivo*, enabling improved epitope targeting and expanding B cell repertoire diversity ^29,30^. Despite promising results in preclinical studies and early-phase clinical trials for viral pathogens ^22,24,31,32^, computationally designed nanoparticle vaccines have yet to be successfully applied to bacterial pathogens. A prominent obstacle is the structural and functional complexity of bacterial antigens, and the limited availability of epitope landscapes at high-resolution, which combined represents a significant challenge to perform rational vaccine design against bacterial pathogens.

In this study, we used deep learning tools including ProteinMPNN ^33^ and AlphaFold2 ^34^ to *de novo* design a new epitope construct that structurally mimics a protective epitope previously identified in domain 3 of SLO ^35^. The epitope-fusion construct was mixed with the assembling component *in vitro* to form an icosahedral symmetry protein-based nanoparticle, a first-in-class, structure-based, epitope-centric SLO-based nanoparticle vaccine against GAS. Immunisation with immunogen resulted in enhanced and higher epitope-specific IgG titres, compared to the community-standard detoxified SLO vaccine construct. Our results demonstrate the importance of constructing epitope-centric immunogens to direct the antibody response towards conserved and protective epitopes of significant biological relevance.

## Results

### Epitope design and assembly of the epitope-centric protein nanoparticle immunogen

In previous work, we used a neutralising streptolysin O (SLO) monoclonal antibody and multimodal protein mass spectrometry (MS) to perform antibody-guided identification of a protective epitope located in domain 3 (D3) ^35^. The epitope is comprised of two discontinuous regions in D3, separated by an elongated subsequence of domain 1 (D1) (**Fig. 1a**). To enable recombinant production of a correctly folded epitope structure, we iteratively applied a *de novo* generative tool, ProteinMPNN, to replace the interspersed D1 subsequence with short linkers comprised of different four-residue combinations. Each candidate construct was structurally modelled using AlphaFold2 and stability was simulated using GROMACS. Among these candidates, the design with an Ala-Pro-Asn-Gly four-residue linker region (modified domain 3, D3m) (**Fig. 1a**) was predicted to fold correctly while preserving the structural integrity of the epitope similar to the structure found in native wild-type SLO (wtSLO). Notably, D3m completely preserves the structure of the identified epitope, which is nearly 100% conserved across over 2000 sequenced GAS genomes ^35^.

**Figure 1.**
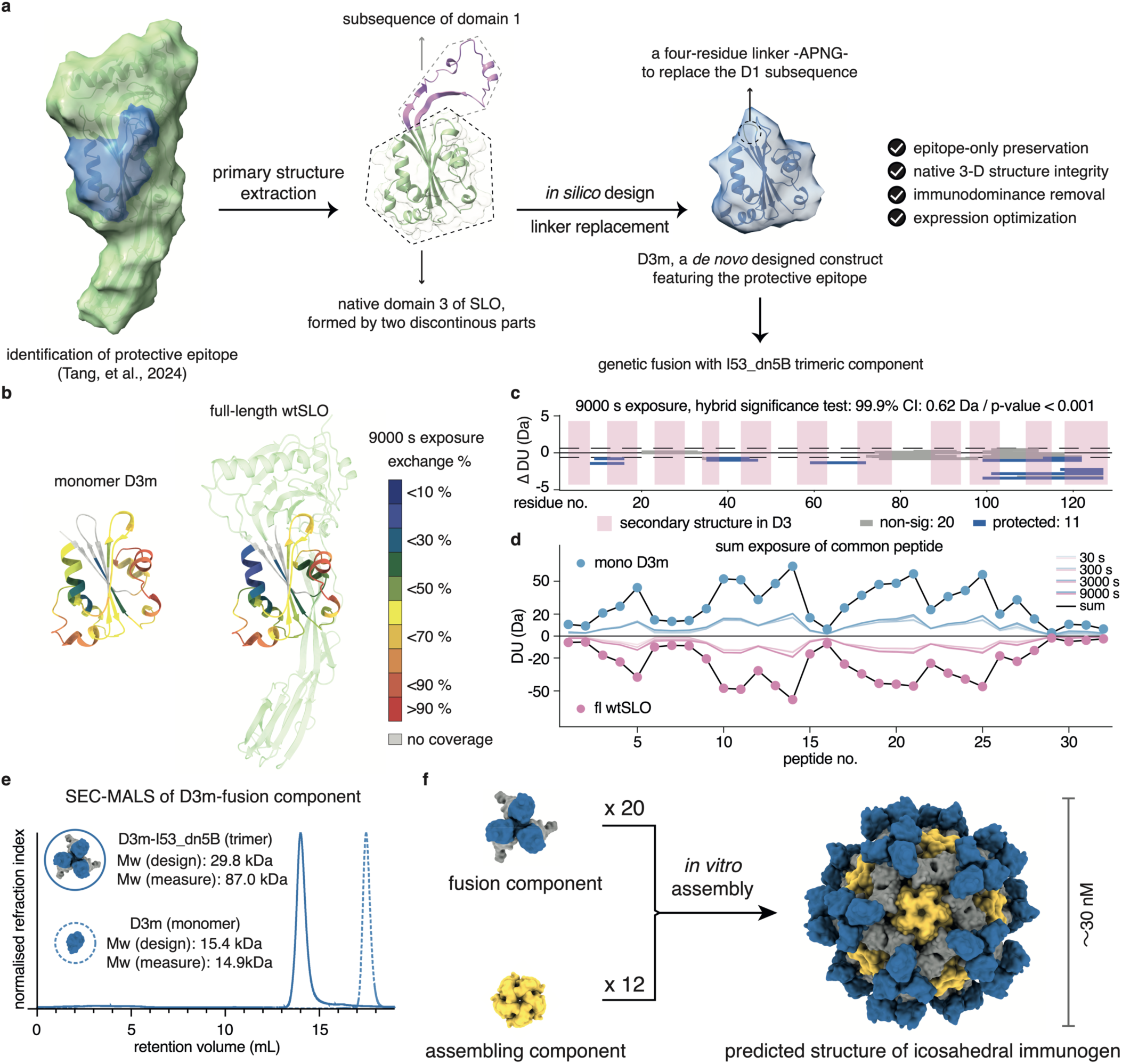
*De novo* design and characterisation of the D3m construct. **a)** Overview of the protein design strategy for the novel immunogen construct (D3m), highlighting the protective epitope in blue that was mapped previously by multimodal mass spectrometry. The region coloured blue in SLO structure represents the identified epitope, which was extracted and redesigned. **b)** Hydrogen/deuterium exchange mass spectrometry (HDX-MS) was used to compare the protein dynamics of the domain 3 region between the D3m construct and the full-length wild-type streptolysin O (wtSLO). Peptide deuteration percentages are colour-coded on the corresponding structures. **c)** A Woods plot showing the differential deuterium uptake of common peptides identified when comparing the full-length wtSLO protein to the monomeric D3m construct. The alpha-helix or beta-sheet motifs are highlighted by pink frames. **d)** A double Butterfly plot summarising both the individual and cumulative deuterium uptake of peptides identified in both the monomeric D3m and full-length wtSLO proteins. **e)** Size-exclusion chromatography coupled with multi-angle light scattering (SEC-MALS) analysis of the monomeric D3m and D3m-fused trimer constructs. The solid line represents the trimeric D3m-fusion, and the dotted line represents the monomeric D3m. Molecular weight (Mw) values are indicated. **f)** Schematic representation of the assembly of the D3m-I53_dn5 nanoparticle (or D3m-NP), comprised of 20 copies of the D3m-I53_dn5B trimer and 12 copies of I53_dn5A pentamer. The D3m construct shown in blue and I53_dn5B in grey. Each nanoparticle displays 60 copies of the D3m construct (coloured in blue).

The new 125-residue D3m epitope was then produced in recombinant form. To verify the solution-state properties of D3m, we employed hydrogen/deuterium exchange mass spectrometry (HDX-MS) to compare the protein dynamics and solvent accessibility between D3m and native full-length wtSLO, focusing on shared domain 3 sequence (**Fig. 1b-d**). Overall, the HDX-MS data demonstrated that D3m exhibits largely the same conformation, stability, and structural order of motifs compared to the native D3, although a slight destabilisation (*i.e.* relative deprotection) was observed in the N- and C-terminal helices in the D3m construct compared to native D3 (**Fig. 1c**). Given the relatively small size of D3m, we hypothesised that presenting D3m on a two-component computationally designed self-assembling protein nanoparticle with icosahedral symmetry (I53_dn5) would enhance immunogenicity ^28^. To achieve this, D3m was genetically fused to the trimeric component I53_dn5B to generate an epitope loaded fusion component (D3m-I53_dn5B), which was recombinantly expressed and purified. Size-exclusion chromatography coupled with multi-angle light scattering (SEC-MALS) of D3m and D3m-I53_dn5B revealed two peaks with the expected molecular weights (**Fig. 1e**). Next, the nanoparticle was *in vitro* assembled by mixing the fusion component (D3m-I53_dn5B) with the assembly (I53_dn5A), to generate an epitope-loaded nanoparticle denoted as D3m-I53_dn5 (or D3m-NP) (**Fig. 1f**). In addition, we prepared and assembled a nanoparticle construct without any epitope, denoted as I53-dn5.

### Structural characterisation of the epitope-centric protein nanoparticle immunogen

SEC-MALS of the two *in vitro* assembled nanoparticle constructs (D3m-NP and I53-dn5) revealed two predominant peaks with molecular weight measurement corresponding to the theoretical molecular weights of the two icosahedral protein nanoparticles with/without D3m (**Fig. 2a**). Native-PAGE and negative-stain electron microscopy (NS-EM) further confirmed the higher-order assembly, homogeneity, and mono-dispersity of both the unmodified I53_dn5 nanoparticle and the D3m-NP displaying D3m immunogens (**Extended Data Fig. 1a-b**).

**Figure 2.**
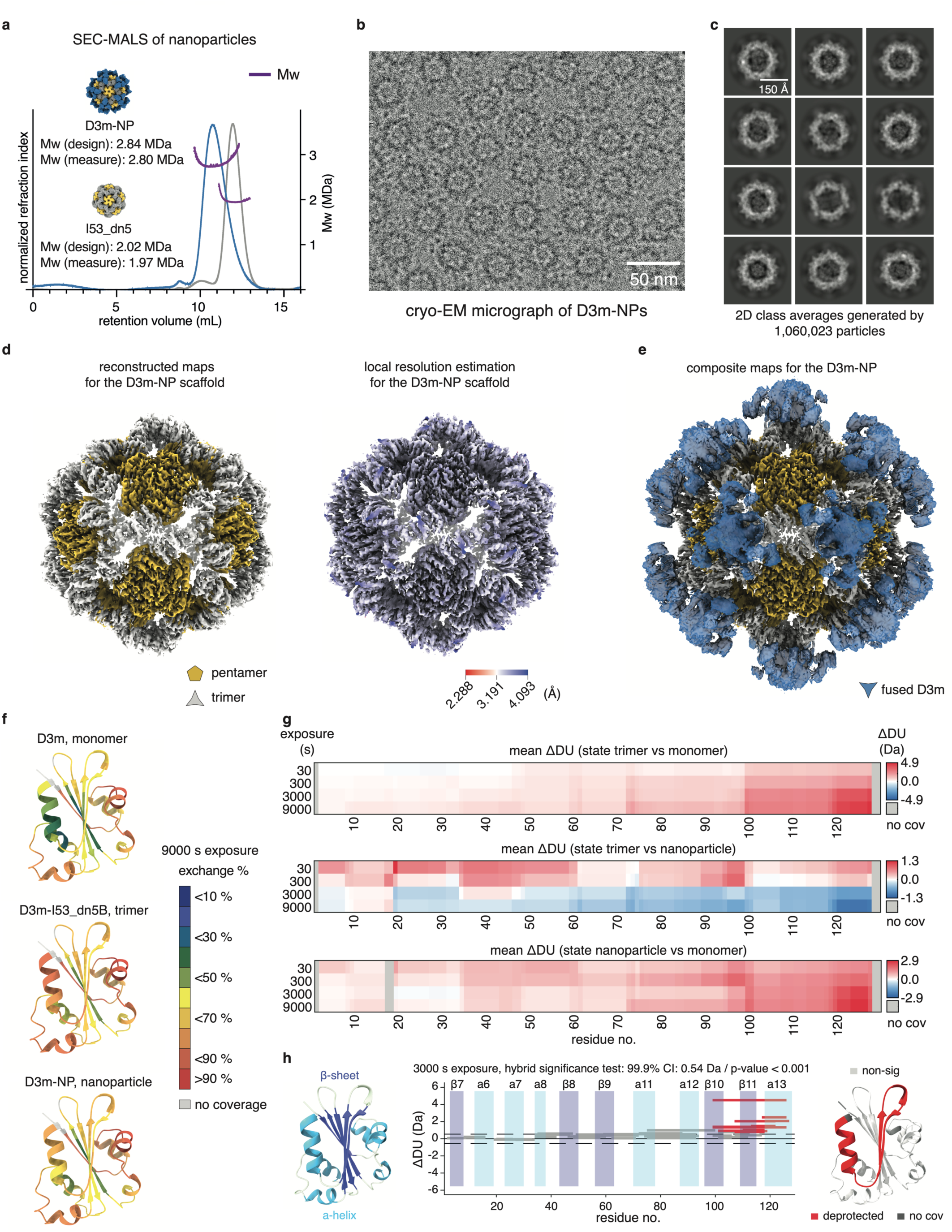
Biophysical characterisation of the assembled protein nanoparticle D3m-NP. **a)** SEC-MALS analysis of both the D3m-NP (blue line) and the control nanoparticle I53_dn5 (grey line) that does not display the epitope. Mw measurements are shown as a purple dotted line. **b)** A representative cryogenic electron micrograph of D3m-NP nanoparticles acquired on a Glacios 200 kV cryo-transmission electron microscope. Scale bar: 50 nm. **c)** 2D class averages obtained from single-particle cryo-EM analysis. Scale bar: 15 nm. **d)** Reconstruction map of the scaffold region from D3m-NP particles. Right: each building block is coloured separately. Left: the colour gradient indicates the estimated local resolution. **e)** Composite maps of D3m-NP after iterative local refinement. No model fitting was applied to the surface-displayed immunogens. **f)** HDX-MS was used to compare the protein dynamics of the D3m immunogen in i) the monomeric state, ii) the trimeric state, and iii) the nanoparticle-presenting state. Peptide deuteration percentages are colour-coded separately on the D3m structure. **g)** A panel of barcode plots demonstrating the mean differential change in deuterium uptake of common D3m peptides among multiple comparisons (i. trimer to monomer, ii. trimer to nanoparticle and iii. nanoparticle to monomer). The deuteriation experiment was carried out using labelling intervals of 30, 300, 3000 and 9000 s. **h)** A D3m structure coloured by the secondary structures. A Woods plot showing the D3m-derived peptides with significant changes in deuterium uptake in the nanoparticle state compared to monomeric state. All HDX-determined peptides were then projected onto the D3m structure and coloured based on significant deprotection.

To determine the structure of the nanoparticle immunogen, D3m-NP was vitrified, and cryo-electron microscopy (cryo-EM) was used to reconstruct the D3m-NP construct. A representative cryogenic electron micrograph showed that the construct formation did not aggregate (**Fig. 2b**). In the initial single particle analysis and subsequent image averaging, D3m was poorly aligned, suggesting that the four-mer linker connecting the D3m immunogen to the trimeric fusion component introduces flexibility (**Extended Data Fig. 3a**, **Fig. 2c**). A cryo-EM *ab initio* reconstruction followed by global refinement yielded a 2.98 Å resolution map in which the structure of two-component scaffold was well defined (**Fig. 2d**). To better visualise the displayed D3m immunogen, local refinement was performed (**Extended Data Fig. 3a**), resulting in an improved reconstruction that revealed the occupancy of D3m at the nanoparticle vertices. The cryo-EM data confirm that D3m-NP displays exactly 60 copies of the D3m immunogen and that the genetic fusion of D3m did not disrupt the intended co-assembled icosahedral architecture (**Fig. 2e**). These results are in line with previously reported reconstruction maps of both unmodified and different antigen-loaded nanoparticles ^23,36^.

Next, we applied HDX-MS to evaluate the conformation and to monitor the protein dynamics of the D3m epitope presented in the monomer, the fusion trimer, and the assembled nanoparticle. On average, more than 96.6 % sequence coverage of overlapping peptides was achieved, which enabled differential analysis of the deuterium uptake and temporal stability across the three constructs (**Extended Data Fig. 2b, Fig. 2f** and **Fig. 2g**). Overall, the three constructs were highly similar over the first 100 residues. The monomer construct was the most protected, followed by the nanoparticle construct, while the trimer construct was the most deprotected. Prominent relative deprotection was, however, observed near the C-terminus in the assembled nanoparticle, which is the region connected to the trimeric fusion component I53_dn5B. Domain 3 of SLO is comprised of 11 secondary structure motifs including six α-helix (α6-8, α11-13) and five β-sheet (β7-11). To pinpoint the motifs with the largest differences, we performed a hybrid significance test. This analysis revealed that the β11 and α13 regions of D3m-NP were most significantly deprotected compared to monomer D3m (**Fig. 2h**). Collectively, the conducted biophysical analysis confirms the successful construction of an SLO D3m-centric protein nanoparticle and the preparation of a monodisperse, non-aggregating vaccine candidate, with the solution-state properties of the D3m-NP-scaffolded epitope closely resembling those in the monomeric form.

### Epitope-centric protein nanoparticle immunogen elicited a potent antibody response

We next compared to what extent the D3m monomer, full-length detoxified SLO (dSLO), and D3m-NP could elicit antibody responses against domain 3 in C57BL/6J mice. The protein content for each construct was normalised at the molar level where in total 2 µg of domain 3 was used per mouse for both the D3m monomer and D3m-NP. In contrast, 10 µg of detoxified SLO was used in accordance with previous studies ^37,38^. All constructs were formulated with adjuvant AddaVax. Four groups with five mice per group were immunised with three doses of the corresponding immunogens plus adjuvant or adjuvant alone (PBS as control group) at two-week intervals, and blood sampling (for serum) was performed at specified intervals (**Fig. 3a**).

**Figure 3.**
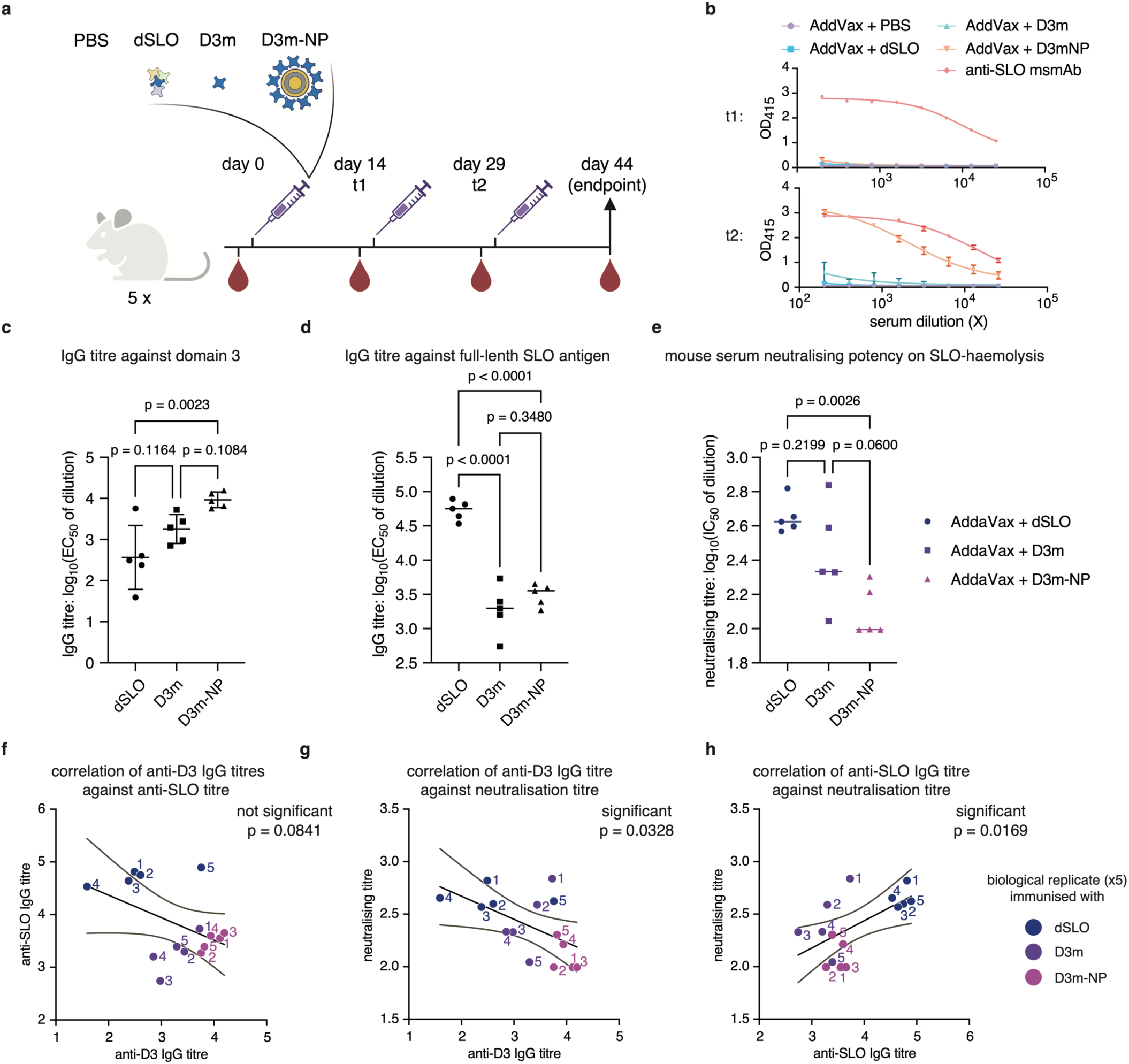
Antibody responses induced by different immunogen constructs. **a)** Schematic overview of the design of the immunisation cohort. dSLO: detoxified full-length SLO immunisation; D3m: D3m monomer immunisation; D3m-NP: D3m-nanoparticle immunisation. PBS group was included as negative control. **b)** The connected dot lines show ELISA measurement of domain 3-specific polyclonal IgG titres across the immunisation groups after the first and second dose. PBS group and anti-SLO mouse antibody was included as negative and positive controls. **c)** Endpoint titres of domain 3-specific antibodies and **d**) endpoint titres of full-length SLO-specific binding IgG antibodies. Each dot represents serum from an individual animal, and the IgG titre is defined as the log_10_-transformation of the dilution factor required to reach half-maximum binding to the corresponding antigen (D3m or dSLO). An ANOVA test was applied for multiple comparisons. The adjusted P-values are displayed on the top. **d)** Quantification of serum antibody neutralising titres induced by each immunogen. Each dot represents serum from an individual animal, and the neutralising titre is defined as the log_10_-transformation of the dilution factor required to reach half-maximum inhibition of SLO-mediated haemolysis. An ANOVA test was applied for multiple comparisons. The adjusted P-values are displayed on the top. **f**-**h**) Linear regression analysis was used to determine the correlation between **e**) D3-specific titre and SLO-specific titres; **f**) D3-specific titres and neutralising serum antibody titres; and **g**) SLO-specific titre and neutralising serum antibody titres. Each dot represents an individual mouse from one of the immunisation groups, with five biological replicates per group.

In this cohort, no antibody titres were detected against D3m or full-length dSLO in the control group that received only adjuvant. In contrast, sera harvested from the other three immunisation groups contained specific IgG antibodies that recognised both the D3m immunogen (**Fig. 3b**-**c**) and the native D3 of SLO (**Fig. 3d**). Although the same protein amount of D3m was administered for immunisation, D3m-NP elicited a significantly >100-fold higher titres already after two immunisation doses compared to both D3m and dSLO (**Fig. 3b**). Furthermore, sera from all three immunisation groups protected against SLO-induced haemolysis (**Fig. 3e**). To determine the correlation between titres and neutralisation, we performed linear regression analysis. Here, the levels of anti-D3 IgG titre, anti-full-length SLO IgG titre, and serum neutralising titres was compared for the three constructs (**Fig. 3f**-**h**). This analysis demonstrates that the D3m-NP immunogen elicited a potent epitope-centric and consistent antibody response against D3 compared to D3m and SLO. However, the contribution of antibodies directed against other epitopes in dSLO improved the overall inhibition against haemolysis. These results indicate that presenting bacterial epitopes in an ordered array on nanometre-scale icosahedral nanoparticles effectively enhance the antibody response towards the displayed epitope. The results furthermore imply that there are other epitopes in SLO that are more relevant for neutralising the cytolytic function of SLO or that combinations of multiple epitopes from SLO is required for maximum inhibition of cytolysis. In summary, we present a first-in-class, structure-based, epitope-centric nanoparticle vaccine against GAS, utilising an icosahedral symmetry protein-based scaffold. Two-component, co-assembling nanoparticle platform applied here enables the selective and modular display of a biologically relevant epitope derived from GAS, eliciting robust and epitope-focused antibody responses. In future studies, antibody-informed combination of multiple epitopes, either from the same antigen or from distinct antigens, provides a promising strategy for the rational design and development of multicomponent, multivalent, epitope-targeted immunogens against GAS.

## Discussion

In this study, we developed a protein-based nanoparticle immunogen to elicit epitope-focused antibody responses against SLO in a mouse model. Our results highlight the benefit of rationally designed immunogens to focus the adaptive immunity response towards conserved and protective epitopes of biological relevance. The strategy encompasses *de novo* epitope design integrated with an emerging vaccine platform. The ability to precisely engineer antigen or epitope structures on designed nanoparticles offers an alternative solution to current state-of-the-art GAS vaccine development programs.

Over the past decades, numerous studies have successfully defined and characterised important GAS virulence factors, including both M-proteins and non-M proteins, as promising antigens for vaccine design ^4,15,39,40^. However, a few obstacles remain unsolved such as the imprecise characterisation of protective epitopes, particularly conformational epitopes related to effective B-cell immunity. Such information is intrinsically challenging to retrieve using conventional epitope mapping methods like peptide scanning arrays and alanine scanning mutagenesis ^41^. Another challenge is the deciphering of the functional correlates between B-cell and T-cell epitopes. The use of toxoids or truncated bacterial antigens with unknown solution-state conformations and with the risk of aggregation at high concentrations could result in the triggering of insufficient or even incorrect immune responses ^42,43^. This can be particularly problematic given the immunogenicity hierarchy effects associated with bacterial immunity, where immunodominant sites often lead to preferential epitope targeting and to subsequent ineffective/non-functional antibody responses ^20^.

The D3m-NP did induce robust anti-D3 IgG titres as compared to monomer D3m and detoxified SLO. The high titres, however, did not directly translate into an equivalent increase in the inhibition of cytolysis. These results suggest that the design of the fusion trimer could be further improved by altering for example the conformation ^44^ or the rigidity of the scaffolded D3m ^45^. Subtle changes made to the epitope or fusion component such as more flexible linkers or rigid spacer repeat sequences that modify the surface exposure or structural stability, may broaden the elicited antibody response after immunisation. In future work, profiling the cellular and plasma proteome can be performed to decipher the molecular signatures and biological processes associated with the different SLO-based constructs included in this study. Such information has the potential to pinpoint immune profiles induced after immunisation. In parallel, given that SLO, like other cholesterol-dependent cytolysins ^46^, harbours T-cell epitopes, additional studies are needed to investigate the T-cell responses activated by these constructs to determine to what extent cellular immunity is influenced by the epitope-centric design and the nanoparticle delivery system ^29,30,47^. Furthermore, GAS infection challenge models are required to assess if the various SLO-based vaccine candidates can mediate protection *in vivo*.

This study demonstrates that the combination of emerging deep learning tools with versatile analytical techniques empowered by the rapid development of protein mass spectrometry have the potential to advance the rational design for next-generation GAS vaccines. Understanding the native protein dynamics of antigens in unbound and in antigen-antibody complex states can elucidate the structural insights of optimal B-cell epitopes that are essential for immune cell recognition and biological functionality. The use of multimodal mass spectrometry can under near-physiological conditions pinpoint important epitopes with sufficient resolution in a relatively high-throughput manner ^35,48–51^. In addition, the accelerating field of immunopeptidomics provides new opportunities to identify linear peptide epitopes of relevance for T-cell immunity ^52,53^. Such advances can be harnessed to further investigate the functional alignment between B-cell and T-cell epitopes in mounting a protective and long-lasting immune response. Emerging vaccine modalities, such as the two-component self-assembling nanoparticle system used in this study, offer further opportunities to present multiple epitope or antigen structures within a single particle, which has the potential to enhance antigen uptake and exhibits favourable pharmacokinetics and biodegradable properties ^47^.

## Methods

### Protein design and production

The full-length streptolysin O (SLO) toxin and its detoxified variant (toxoid), derived from the UniProt ID: P0DF97 sequence, were produced as previously described ^35,54,55^ . Briefly, the detoxified SLO (dSLO) was generated by substituting both Thr_564_ and Leu_565_ with glycine residues to abolish haemolytic activity. An N-terminal composite affinity tag (*i.e.* Strep II-hemagglutinin-histidine-tobacco with virus protease recognition site) was fused to both sequences. Plasmids encoding tagged wild-type SLO (wtSLO) and dSLO were synthesised and assembled by the Lund University Protein Production Platform (LP3). These plasmids were transformed into *E. coli* BL21(DE3) competent cells (Thermo Fisher Scientific). Recombinant protein production followed standard protocols, using 0.1 mM IPTG (Thermo Fisher Scientific) for inducing, BugBuster (Novagen) for cell lysis, and Ni^2+^-charged immobilised metal affinity chromatography (IMAC) resin (Bio-Rad) for two-step gravity-flow purification.

The immunogen, D3m, was designed to feature a previously determined protective epitope supported by two discontinuous regions of SLO (residues 250-299 and 346-420). Using ProteinMPNN ^33^ and AlphaFold2-Multimer ^34,56^, the interrupting subsequence was replaced with a short four-residue linker (-Ala-Pro-Asn-Gly-), resulting in a compact construct intended to mimic the native conformation of domain 3 in wtSLO. To facilitate nanoparticle assembly, the I53_dn5B trimeric component was genetically fused to the C-terminus of D3m via a flexible linker (-Gly-Gly-Ser-Ala-). All constructs (D3m, D3m-I53_dn5B, and I53_dn5A) included a C-terminal His-tag for purification. Protein synthesis, purification, and characterisation were performed by Protein Production Sweden (PPS) at Umeå University. Protein purity and sequence integrity were verified using SDS-PAGE (Bio-Rad) and bottom-up mass spectrometry. Samples were stored at -80 °C until further use.

### Nanoparticle assembling and biophysical characterisation

SEC-MALS was used to assess the homogeneity and oligomeric state of various components and assembled nanoparticles. Lipopolysaccharide (LPS) was removed from all building blocks and dSLO, using endotoxin removal spin columns (Thermo Fisher Scientific). Nanoparticle assembly was performed *in vitro* by mixing I53_dn5A with either D3m-I53_dn5B or unmodified I53_dn5B at an equimolar subunit ratio. The mixture was incubated for 10 minutes at room temperature (RT) with agitation at 1,000 rpm. The mobile phase for all D3m-derived constructs was buffer-exchanged to 25 mM Tris-HCl, pH 8.0, 500 mM NaCl, and 5% v/v glycerol.

Approximately 50 µL of each sample, at a concentration of ∼2 mg/mL, was injected in triplicate into an OMNISEC system (Malvern Panalytical). The system included the OMNISEC RESOLVE module, equipped with either a Superdex 200 Increase 10/300 GL or Superose 6 Increase 10/300 GL column (Cytiva), and the OMNISEC REVEAL, an integrated multi-detector module comprising right-angle light scattering (RALS) at 90°, low-angle light scattering (LALS) at 7°, a differential refractive index detector, a viscometer, and a diode-array-based UV/VIS spectrometer. Bovine serum albumin (BSA; Thermo Fisher Scientific) was used to normalise detector responses. Data were collected and analysed using OMNISEC v11.41 software, with the flow rate maintained at 0.5 mL/min. Prism 10 software was subsequently used for data plotting and visualisation. Native-PAGE experiment (Thermo Fisher Scientific) was also performed on the building blocks and assembled nanoparticles according to the manufacturer’s instructions.

### Hydrogen/deuterium exchange mass spectrometry and data analysis

The HDX-MS experiments were performed using an automated sample handling system set up with a LEAP H/D-X PAL platform (Trajan Scientific and Medical) interfaced to an LC-MS system comprising an Ultimate 3000 micro liquid chromatography unit and an Orbitrap Q Exactive Plus mass spectrometer (Thermo Fisher Scientific). The HDX experiments were conducted on wtSLO, dSLO, D3m, D3m-I53_dn5B, and D3m-NP protein constructs. Protein labelling with D_2_O was carried out at intervals of 0, 30, 300, 3,000, and 9,000 seconds at 4 °C, followed by quenching with a solution containing 1% TFA, 0.4 M TCEP, and 4 M urea, also at 4 °C. The quenched samples were immediately injected for on-line digestion using a mixed protease column of Nepenthesin-2 and pepsin.

The digested peptides were subjected to on-line solid-phase extraction and washing with 0.1% formic acid for 60 seconds on a trap column (PepMap300 C18, Thermo Fisher Scientific). Subsequently, the trap column was switched in-line with a reversed-phase analytical column (Hypersil GOLD, Thermo Fisher Scientific). Peptide separation was performed at 1 °C using mobile phases of 0.1% formic acid (solvent A) and 95% acetonitrile with 0.1% formic acid (solvent B). A gradient of 5 to 50% solvent B was applied over 8 minutes, followed by an increase from 50% to 90% solvent B over 5 minutes. MS analyser was equipped with a heated electrospray ionisation (HESI) source. The capillary temperature was set at 250 °C, with sheath gas, auxiliary gas, and sweep gas flow rates set at 12, 2, and 1 arbitrary unit, respectively. Full MS scan spectra were acquired at 70,000 resolution, with an automatic gain control target of 3e^6^, a maximum ion injection time of 200 ms, and a scan range of 300 to 2,000 m/z. Identification of digested peptides was achieved by analysing un-deuterated control samples using data-dependent acquisition (DDA) tandem MS. A pooled peptide library, including sequence, charge state, and retention time, was generated by running mixed protease-digested, un-deuterated samples against the wtSLO and D3m sequences using PEAKS Studio X (Bioinformatics Solutions Inc.).

HDX data analysis and visualisation were performed using HDExaminer v3.4.2 (Sierra Analytics Inc.). Deuterium incorporation for identified peptides was calculated as the relative mass difference between deuterated and non-deuterated peptides without applying back-exchange correction using fully deuterated samples as references. Each spectrum was manually inspected, and peptides with low scores, significant outliers, or inconsistent retention times were excluded from further analysis. To compare different constructs, individual charge states for each peptide were combined. The differential deuterium uptake data were further analysed using Deuteros 2.0 ^57^ to perform hybrid significance testing and visualise results as Woods plots, butterfly plots, and barcode plots. This combination of analytical tools enabled detailed multiple comparative analysis of the structural dynamics and solvent accessibility among the various constructs.

### Negative-stain electron microscopy and cryo-EM of the nanoparticle immunogens

For imaging nanoparticles with or without immunogen loading, D3m-NP and NP samples were diluted to 0.05 mg/mL and buffer-exchanged into 25 mM Tris, pH 8.0, 150 mM NaCl. Grids were glow-discharged immediately prior to use. A 3 µL aliquot of the sample was applied to the grid and incubated for 1 minute before blotting away excess liquid. Grids were stained with 3 µL of 0.75% (w/v) uranyl formate, and the excess stain was immediately blotted away. Images were acquired on a Talos L120C 120 kV electron microscope.

For cryo-electron microscopy (cryo-EM) reconstruction of the D3m-NP nanoparticle, 3 µL of 2.8 mg/mL D3m-NP was applied to a freshly glow-discharged holey carbon grid (1.2 µm hole, 1.3 µm spacing, 200-mesh). Multiple blotting steps were used before plunge freezing with a Vitrobot Mark IV (Thermo Fisher Scientific) under 100% humidity at 4 °C, using a blot force of 0 and a blotting time of 2-4 seconds. Screening data were collected on a Glacios 200 kV cryo-transmission electron microscope (cryo-TEM) equipped with a Falcon 4i Direct Electron Detector (4k x 4k pixels) and CETA-D 4k x 4k CMOS. The system was operated using EPU data collection software with AFIS mode. The dose rate was set to 10.82 e/Å^2^, and each movie was acquired in counted mode, fractionated into 126 frames with an exposure time of 29.3 ms per frame. Analysing data were further collected on a Titan Krios 300 kV cryo-TEM coupled with a Falcon 4i direct electron detector (4k x 4k pixels) combined with Selectris (energy filter) and CETA 4k x 4k CMOS. The system was operated using EPU data collection software. The dose rate was set to 40 e/Å^2^, and each movie was acquired in counted mode, fractionated into 40 frames with an exposure time of 59.2 ms per frame.

A total of 11,291 micrographs were collected in a single session. Patch contrast-transfer-function parameters, particle picking, and 2D classification were carried out using CryoSPARC v4. After initial screening, 1,060,023 particles of the highest-quality classes were selected and subjected to non-uniform refinement with icosahedral symmetry applied, resulting in a global reconstruction at 2.98 Å resolution. After iterations of local refinement with masks applied, in **Figure 2**, high contour levels (yellow/grey) and low contour levels (blue) of the 3-D reconstruction are combined to a composite map to visualise the I53_dn5 scaffold and the displayed 20 D3m-trimer (*i.e.* 60 copies of D3m immunogen loaded per particle), respectively. Reported resolutions are based on the gold-standard Fourier Shell Correlation (FSC) criterion at a threshold of 0.143.

### Mouse immunisation study

All animal procedures were conducted in compliance with ethical guidelines and approved by the Malmö/Lund Institutional Animal Care and Use Committee under ethical permit number 11542-2020 and 17747-2024. Female 6-week-old C57BL/6J mice (Janvier) were anaesthetised with isoflurane and injected intramuscularly into the right flank with one of the following: PBS, detoxified SLO (10 µg), SLO-D3m (2 µg), or SLO-D3mNP (6 µg). Each immunogen was formulated with adjuvant (AddaVax, InvivoGen) in a 1:1 volume ratio, resulting in a total injection volume of 50 µL. Injections were administered using a 27 G needle (BD Microlance 3) on days 0, 14, and 29 accordingly. Blood samples for serum preparation were collected via tail vein on days -1, 13, and 28. On day 44, animals were sacrificed, and blood was obtained by cardiac puncture, and serum was prepared by centrifugation at 2,000 x g for 10 minutes. Serum samples were stored at -20 °C or -80 °C for long-term storage.

### Indirect ELISA and SLO-haemolysis inhibition assay

Antigen-specific IgG titres were measured using an indirect enzyme-linked immunosorbent assay (ELISA). Plates were coated with either 5 µg D3m or 2 µg dSLO antigen per well and incubated overnight at 4 °C. The plates were washed three times with PBS containing 0.05% v/v Tween 20 (PBST) and blocked with PBS containing 2% BSA (Thermo Fisher Scientific) for 30 minutes at 37 °C with shaking at 300 rpm using a Thermomixer (Eppendorf). Reference anti-D3m/SLO monoclonal antibody and immune sera samples were serially diluted tenfold and added to the plates, which were incubated at 37 °C for 90 minutes at 300 rpm. Horseradish peroxidase (HRP)-conjugated goat anti-mouse IgG antibody (Bio-Rad, 1:3000 dilution) was added, followed by incubation at 37 °C for 60 minutes at 300 rpm. Freshly made HRP substrate solution (Bio-Rad) was added to develop the signal, and the reaction was stopped with 250 mM H_2_SO_4_. Absorbance was measured at 415 nm wavelength using a plate reader (BMG LABTECH). The binding affinity (avidity) of immune sera-derived polyclonal antibodies towards the given antigens was inferred as EC_50_, calculated as the dilution factor required for half-maximal reactivity. Negative control sera from control (PBS + AddaVax)-immunised mice showed no observed signal above background levels.

To assess the inhibitory effects of immune sera on SLO-induced haemolysis, defibrinated sheep erythrocytes (Thermo Fisher Scientific) were diluted with PBS to a 4% v/v cell suspension. To this erythrocyte suspension, 20 ng of 2 mM TCEP (Thermo Fisher Scientific)-activated SLO toxin and 2 µL of immune sera were added and incubated for 30 minutes at 37 °C. After incubation, the mixture was added to the diluted erythrocyte suspension and further incubated for another 30 minutes at 37 °C. An anti-SLO monoclonal antibody (6D11 clone, Abcam) was used as a positive control, and samples without SLO addition were used as negative controls. The plates were then centrifuged, and the supernatant was transferred to fresh plates for absorbance measurement at 541 nm wavelength using a microplate reader (BMG LABTECH). Absorbance measurement corresponds to the level of released free haemoglobin, indicative of cell lysis triggered by SLO. The neutralising titre of immune sera against SLO-induced haemolysis was inferred as IC_50_, calculated as the dilution factor required for half-maximal haemolysis inhibition.

## Data Availability

The mass spectrometry proteomics data and cryo-EM data will be deposited to the public repository upon completion of peer-review process.

## Acknowledgements

We thank Dr. Annie Dosey and Dr. Neil P. King for sending us the plasmids including I53_dn5A.2 and I53_dn5B included in this study. We gratefully acknowledge the Swedish National Infrastructure for Biological Mass Spectrometry (BioMS), the SciLifeLab Integrated Structural Biology platform, and the Protein Production Sweden (PPS) for providing facilities and experimental support. The data was collected at the Cryo-EM Swedish National Facility funded by the Knut and Alice Wallenberg, Family Erling Persson and Kempe Foundations, SciLifeLab, Stockholm University and Umeå University. We would also like to thank Dr. Mikael Lindberg, Dr. Celeste Sele, Dr. Sara Sandin, and Dr. Tanvir Shaikh for assistance.

---

**Extended Data Figure 1.**
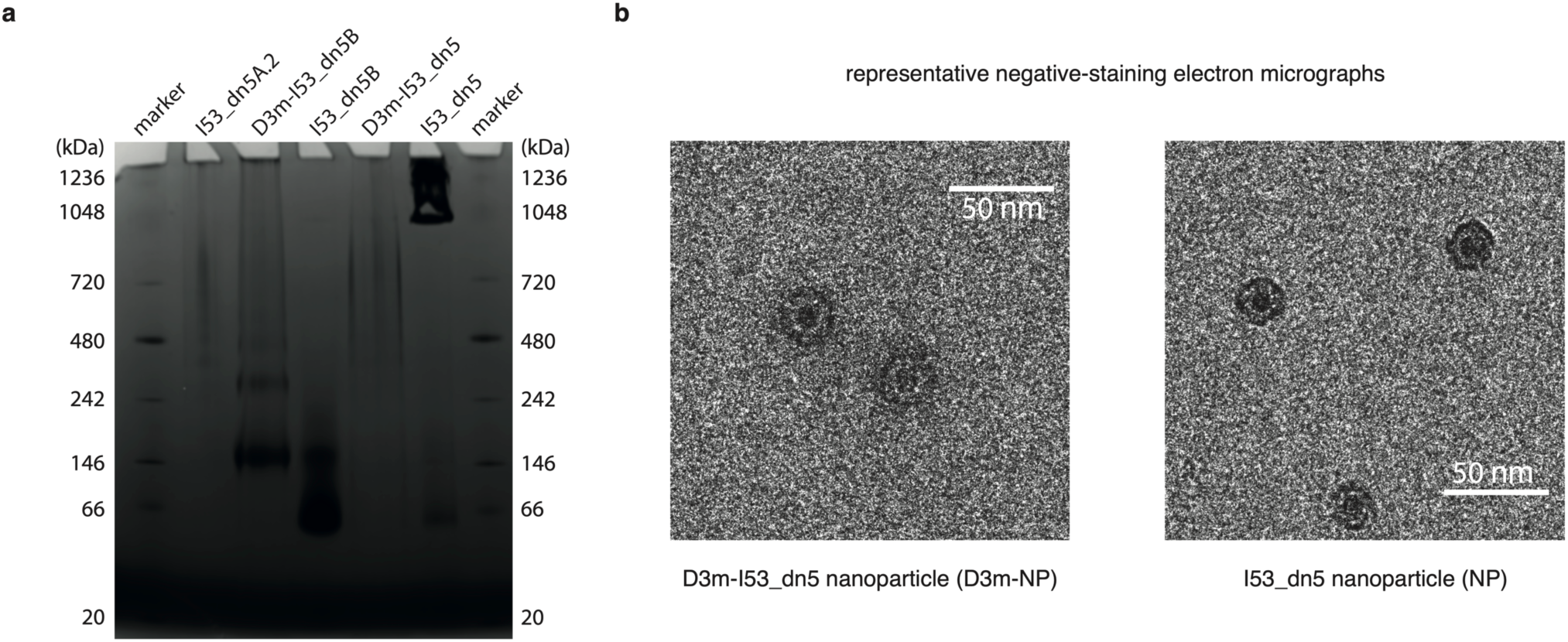
Biophysical characterisation of the assembled protein nanoparticle D3m-NP. **a)** Native-PAGE analysis of the different building blocks separately and assembled into the nanoparticles. **b)** Negative-stain electron microscopy (ns-EM) analysis of assembled D3m-NP nanoparticles. Scale bar: 50nm.

**Extended Data Figure 2.**
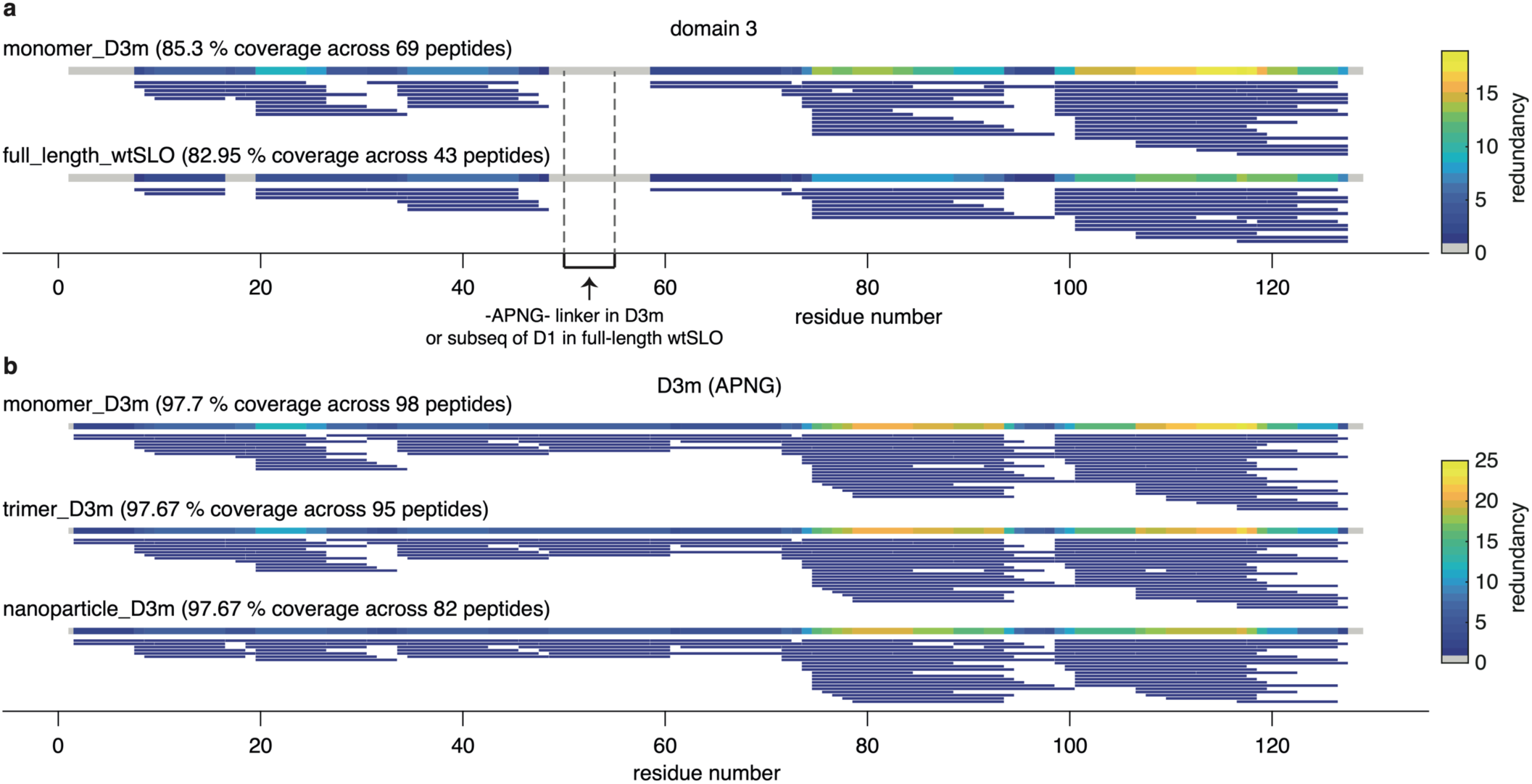
HDX-MS analysis of protein dynamics of different constructs. **a)** Comparison of the peptide coverage and redundancy of overlapped domain 3 sequence between the D3m monomer and full-length wtSLO using HDX-MS. To assess in-solution protein dynamics, D3m monomer and full-length dSLO were deuterated at time intervals of 30, 300, 3000, and 9000 s. Only peptides consistently identified across all time points were included. Differences in the construct sequences are highlighted with dashed lines and annotated accordingly. **b)** Comparison of peptide coverage and redundancy of the D3m region across three different D3m-derived constructs: i) monomer, ii) trimer, and iii) nanoparticle format. To compare in-solution protein dynamics, each construct was deuterated at time intervals of 30, 300, 3000, and 9000 s, respectively. Only peptides consistently identified across all time points were included.

**Extended Data Figure 3.**
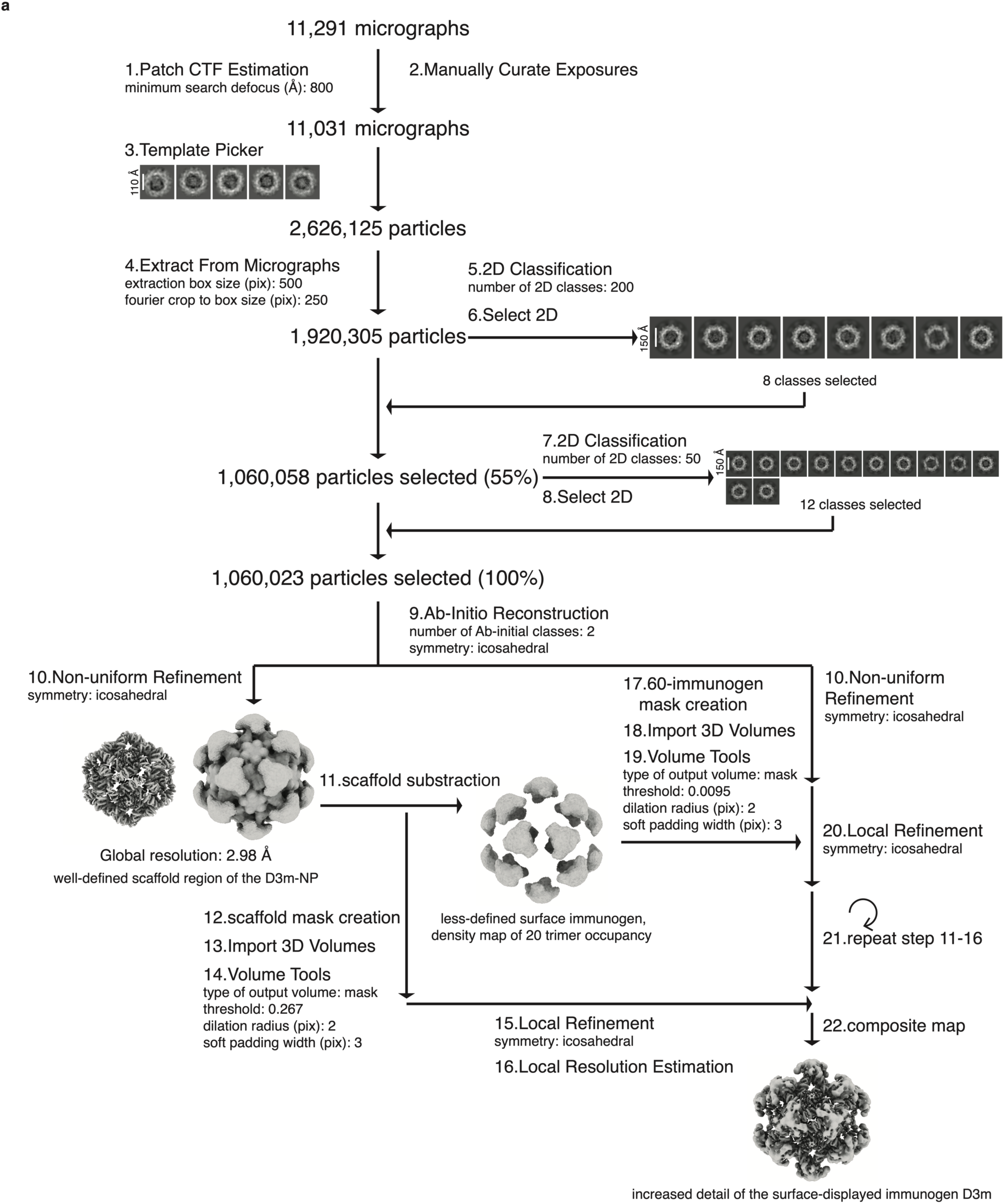
Cryo-EM processing pipeline. **a)** Flowchart showing the image-processing workflow for the cryo-EM dataset of D3m-NP. The process started with 11,291 movies collected on a Titan Krios 300 kV cryo-TEM equipped with a Falcon 4i Direct Electron Detector (4k × 4k pixels) and a CETA 4k × 4k CMOS camera. Contrast transfer function (CTF) estimation for each micrograph was calculated with CryoSPARC v4. Particles were initially manual picked from one micrograph using the CryoSPARC v4 Blob Picker and then sorted by 2D classification using CryoSPARC v4 to assess quality and generate templates. The selected classes from the 2D classification are shown. After particle picking and cleaning by 2D classification, the dataset contained 1,060,023 particles. Particles were used to generate *ab initio* templates in CryoSPARC v4, followed by non-uniform global refinement, and local refinement with different created masks using CryoSPARC v4.

